# Crystal structure of monkeypox H1 phosphatase, an Antiviral Drug Target

**DOI:** 10.1101/2022.09.30.510410

**Authors:** Wen Cui, Haojun Huang, Yinkai Duan, Zhi Luo, Haofeng Wang, Tenan Zhang, Henry Nguyen, Wei Shen, Dan Su, Xiaoyun Ji, Haitao Yang, Wei Wang

**Affiliations:** Institute of Life Sciences, Chongqing Medical University, Chongqing, China; Shanghai Institute for Advanced Immunochemical Studies and School of Life Science and Technology, ShanghaiTech University, Shanghai, China; School of Life Science and Technology, ShanghaiTech University, Shanghai, China; State Key Laboratory of Biotherapy and Cancer Center, West China Hospital, Sichuan University and Collaborative Innovation Center of Biotherapy, Chengdu, China; State Key Laboratory of Pharmaceutical Biotechnology, Department of Biotechnology and Pharmaceutical Sciences, School of Life Sciences, Nanjing University, Nanjing, China; Shanghai Clinical Research and Trial Center, Shanghai, China; Tianjin International Joint Academy of Biotechnology and Medicine, Tianjin, China

## Abstract

The current monkeypox outbreak has caused over 64,000 global cases, but the effective treatments are very limited. The dual specific phosphatase (H1) from monkeypox antagonizes the immune response and is crucial for viral replication, making it an attractive antiviral target. Here we determined a 1.8-Å crystal structure of H1, which forms a domain swapped dimer resembling a butterfly. Each active site, which consists of a Cys-Arg-Asp triad, captures a phosphate ion. The observed conformation mimics the final step of catalysis prior to product release. The crystal structure provides a strong foundation for the discovery of new antivirals against this emerging worldwide pathogen.

---

Since its appearance in May 2022, monkeypox has spread to more than 100 countries and afflicted tens of thousands of people (https://worldhealthorg.shinyapps.io/mpx_global/). Individuals infected with monkeypox present a fever, an extensive characteristic rash, and usually swollen lymph nodes ^1^. The number of confirmed cases worldwide continues to grow at a rapid rate, but the treatment to this highly infectious viral disease is still very limited. Identification of new targeted-therapies will be crucial to control of this emerging public health threat.

The monkeypox virus is an enveloped double-stranded DNA virus that belongs to the *Orthopoxvirus* genus of the *Poxviridae* family^2^. It has a very large genome (∼200 kilo base pairs) and codes around 200 proteins^3^. Poxviruses express a dual specific phosphatase (H1) that de-phosphorylates STAT1 and blocks interferon signal transduction^4,5^. H1 is conserved in poxviruses and is essential for virus replication in cell culture^6^. Inhibiting H1 expression results in greatly reduced infectivity. About 200 copies of H1 are packaged into newly formed viral particles and function in the early stage of viral infection^6^. H1 has also been suggested to de-phosphorylate monkeypox proteins F18, A14 and A17 ^6-8^. Due to the importance of H1 in modulating interferon-signaling and viral replication, it serves as an attractive anti-poxvirus drug target.

To understand the mechanism for monkeypox H1 catalyzed dephosphorylation and provide an accurate structural model for drug discovery, we determined a crystal structure of H1 to 1.8 Å resolution (Fig. 1, Extended Data Table 1). Monkeypox H1 has 171 amino acid residues and the refined model includes residues 2–171 with well-fitting electron density. There is one H1 molecule in an asymmetric unit. Two H1 molecules are related by crystallographic symmetry and form a domain swapped dimer, which resembles a butterfly (Fig. 1a). The overall structure is composed of six α helices and four β strands. A four-stranded β-sheet is sandwiched by helices α2 and α3−α6 on either side. The active site is located near the C-terminus of the last β-strand. The two active sites are ∼39 Å apart (Fig. 1a). There is a phosphate ion captured at each active site, representing the final stage of catalysis before the product is released.

**Fig. 1.**
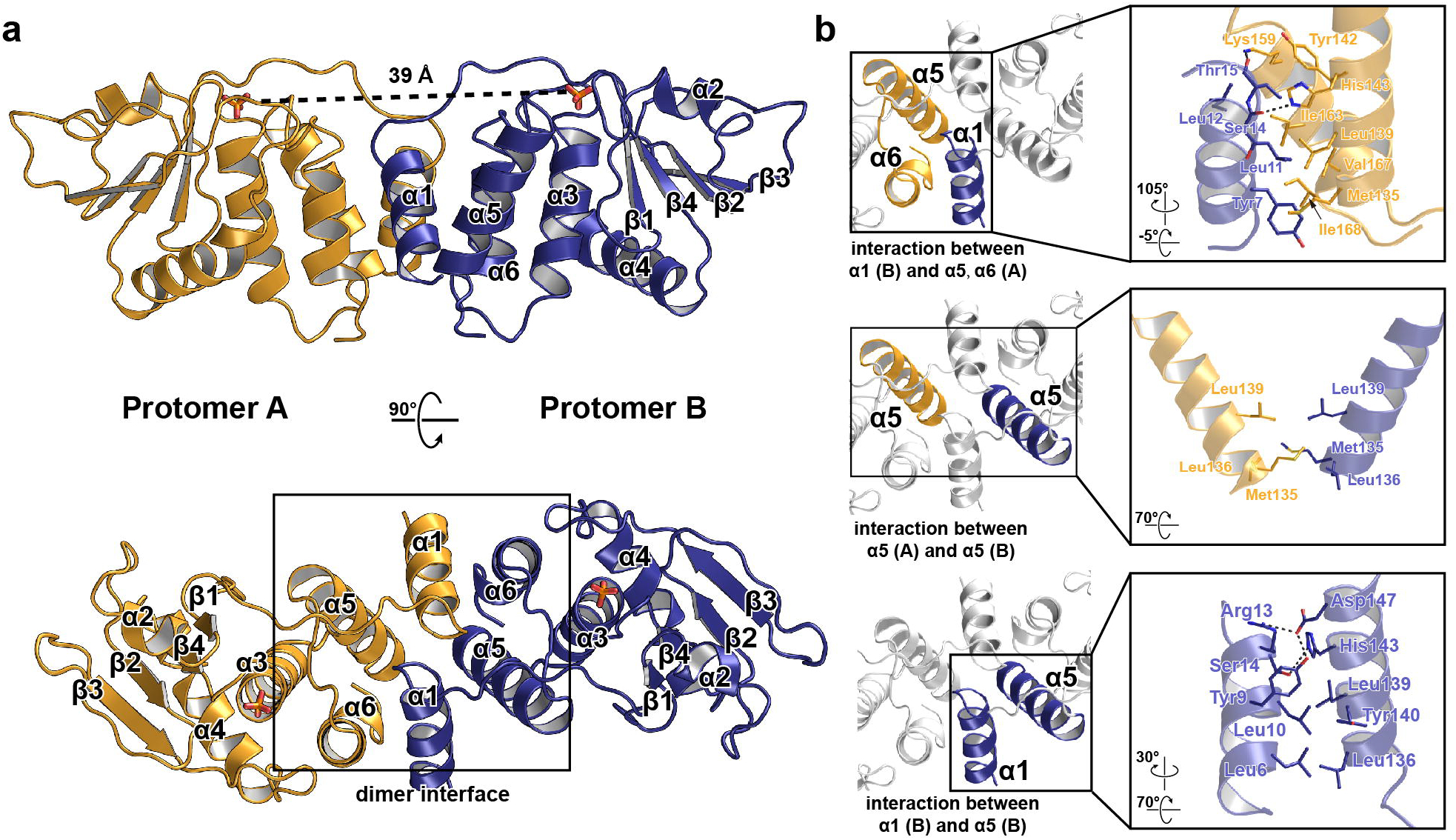
Monkeypox H1 forms a domain swapped dimer. **a**, The overall structure of H1 dimer, shown in two different views. Two protomers are in yellow and blue, respectively. The secondary structure elements are labeled. The two active sites are on the same side of the dimer and ∼39 Å apart. There is a phosphate ion in each active site. **b**, Zoom in of the dimer interface, which consists three components. The residues that participate in dimerization are shown as stick models. Hydrogen bonds are shown as dashed lines.

The N-terminal helix α1 from each protomer interchanges to mediate H1 dimerization. The α1 from one H1 protomer forms a four-helix bundle with three helices α4−α6 of the pairing protomer. The buried surface area between Protomer A and B is ∼1000 Å^2^, which is stabilized by both hydrophilic and hydrophobic interactions. Residues Ser14 and Thr15 in α1 form hydrogen bonds with His143 and Tyr142/Lys159 of the other H1 protomer, respectively; whereas Tyr7, Leu11 and Leu12 participate in hydrophobic interactions with Met135, Leu139, Lys159, Ile163, Val167 and Ile168 from the pairing H1 molecule (Fig. 1b, upper panel). In addition, three residues in α5, Met135, Leu136 and Leu139, face the symmetry related residues in the dimer to expand the hydrophobic interface (Fig. 1b, middle panel). α1 is also stabilized by intramolecular hydrogen bonds and hydrophobic interactions between residues of α1 and α5 (Fig. 1b, lower panel). Size exclusion chromatography confirms the H1 dimer in solution, suggesting that dimerization represents its functional state (Extended Data Fig. 1).

The H1 active site consists of a Cys-Arg-Asp catalytic triad (Fig. 2a). The conserved Cys and Arg residues are in a loop between β4 and α4 (^109^HCVAGVNRS^117^), which is also known as the phosphate-binding loop. The arginine residue (Arg116) of this loop captures the phosphate ion, whose guanidinium group interacts with two phosphate oxygens through two hydrogen bonds with the distances of 2.9 Å and 3.0 Å, respectively (Fig. 2b). This important arginine residue guarantees efficient binding and orientation of the substrate. At the bottom of the catalytic pocket (Fig. 2c), the conserved Cys110 attacks the phosphorous atom during the de-phosphorylation reaction, resulting in a transient enzyme-phosphate intermediate. This intermediate is then hydrolyzed to generate inorganic phosphate and the regenerated enzyme. The sulfur atom of Cys110 is positioned in line with a phosphorous-oxygen (P-O) bond which corresponds to the one formed during the enzyme regeneration step (Fig. 2b). Asp79 is responsible for coordinating the water molecule, which is also hydrogen bonded to an oxygen from the phosphate group (Fig. 2b). This residue functions as a general acid, facilitating both the formation of the enzyme-phosphate intermediate and its hydrolysis. Thus, the crystal structure represents the final step of catalysis before the product is released.

**Fig. 2.**
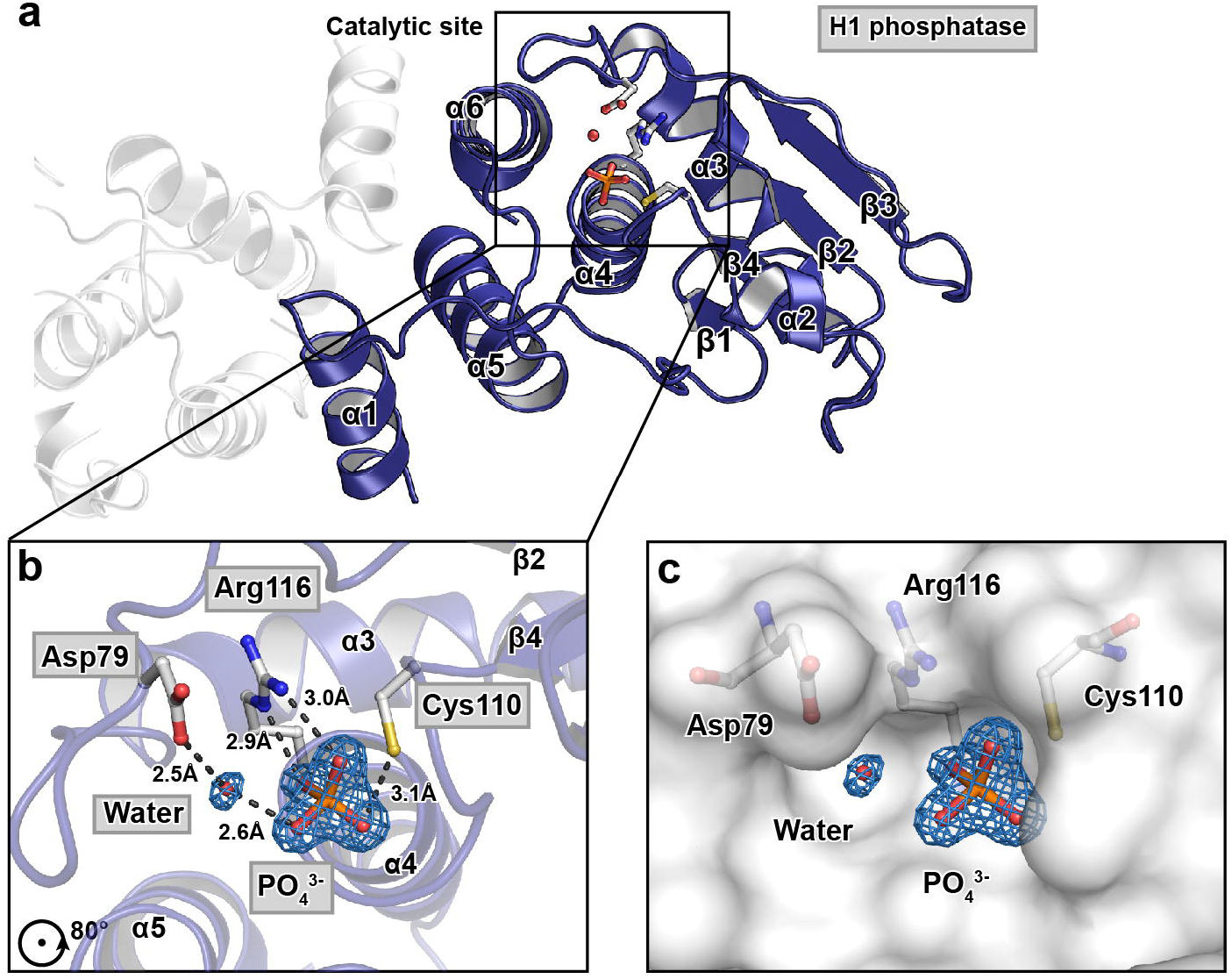
The active center of H1 captures the final step of catalysis before the product is release. **a**, The active site is located near the C-terminus of β4, which consists of a phosphate binding loop (between β4 and α4) and a general acid loop (between β3 and α3). The Cys-Arg-Asp catalytic triad and the phosphate ion in active site are shown as stick models. Water is presented as a red sphere. **b and c**, Zoom in of the active site, where H1 is in cartoon and surface representations, respectively. Hydrogen bonds are shown as dashed lines. The distances between the hydrogen bonded atoms are labeled. Simulated annealing omit map is shown for the phosphate ion and the water molecule. The map is generated with the standard composite omit map procedure implemented in Phenix (torsion angle simulated annealing with 5% of model omitted at a time).

The high-resolution crystal structure of monkeypox H1 reveals at least two hot spots for drug discovery. The first hot spot is the dimer interface, which is unique among the members in the protein tyrosine phosphatase (PTP)/dual specificity phosphatase (DSP) family (Extended Data Fig. 2a). Blocking H1 dimerization may potentially inhibit its ability to dimerize and dephosphorylate the phosphor-tyrosine in activated signal transducer and activator of transcription 1 (STAT1), which is also a homodimer ^9,10^. In addition, the active site is another potential target for inhibition. Although the active sites of all phosphatases in PTP/DSP family are built around a phosphate-binding loop (with a sequence HCX_5_R(S/T)) and have a similar main chain structure, the side chains around the active center are different^10-12^ (Extended Data Fig. 2b), which affects their own substrate specificity and may allow the development of specific inhibitors.

In conclusion, we report a high-resolution crystal structure of the monkeypox H1 phosphatase that lays a solid foundation for its mechanistic study and the discovery of antiviral compounds against this emerging pathogen.

## Methods

### Cloning, protein expression and purification of monkeypox H1

The DNA coding for monkeypox H1 of the current monkeypox virus (MPXV) outbreak (isolate name MPXV_USA_2022_MA001; accession ON563414 in GenBank) was synthesized by Tsingke Biotech (China) and cloned into the pETDuet-1 expression vector using restriction sites *Eco*RI and *Nde*I, coding proteins with a N-terminal His-tag. The plasmid was verified by sequencing (Tsingke Biotech, China).

H1 was expressed in Escherichia coli BL21(DE3) in Luria broth (LB) at 16 °C for 16-18 h with 0.5 mM IPTG. Bacteria expressing H1 were harvested and resuspended in a lysis buffer containing 20 mM Tris-HCl, pH 8.0, 0.5 M NaCl, 10 mM imidazole, 5 mM MgCl_2_ and 10% glycerol and lysed by high pressure homogenization. After centrifugation (16,000 x g, 30 min at 4 °C), the supernatant was loaded onto a Ni-NTA column (GE Healthcare, USA). The column was washed using a buffer containing 20 mM Tris-HCl, pH 8.0, 0.2 M NaCl, 50 mM imidazole, 5 mM MgCl_2_ and 10% glycerol and eluted using a similar buffer supplemented with 250 mM imidazole. The eluted protein was concentrated and purified by gel-filtration chromatography (Superdex 200 10/300 GL, GE Healthcare, USA), using a buffer containing 20 mM Tris-HCl, pH 8.0, 500 mM NaCl and 5 mM MgCl_2_.

### Crystallization, data collection and structure determination

Crystals were grown by the sitting-drop vapor diffusion method. H1 was crystallized at 16 °C by mixing 1 µL protein (15 mg/mL) with 1 µL crystallization buffer containing 0.2 M Li_2_SO_4_, 0.1 M Bis-Tris pH 6.5 and 25% polyethylene glycol 3,350. The crystals were cryoprotected using the crystallization buffer supplied with 20% glycerol. X-ray diffraction data were collected on beamline BL18U1 at the Shanghai Synchrotron Radiation Facility at 100 K and at a wavelength of 0.97776 Å. Data integration and scaling were performed using XDS ^13^. The structure was determined by molecular replacement method with the AutoSol program in PHENIX ^14^ using the structure of Vaccinia Virus H1 (PDB: 3CM3) as a search model. The H1 model was initially built by the Autobuild program in PHENIX and subsequently subjected to iterative cycles of manual building in Coot ^15^ and refinement in PHENIX.

### Structural comparison with human phosphatases in the PTP/DSP family

The coordinates of monkeypox H1 was uploaded on DALI server ^16^ and search for proteins with similar structure in Protein Data Bank (using the PDB50 subset). 30 phosphatase structures were found to have a Z-score higher than 13.0, which indicates significant structural similarity. Among them, two human phosphatases are crystalized as dimer: human dual-specificity phosphatase 5 (hDSP5, PDB ID: 2G6Z) and human dual-secificity phosphatase 27 (hDSP27, PDB ID: 2Y96). Their dimerization mode was compared.

## Data availability

Coordinates and structure factors for Monkeypox virus H1 has been deposited in Protein Data Bank (PDB) with accession number 8GZ4.

## Acknowledgements

We are grateful to the staff at the BL18U1 at Shanghai Synchrotron Radiation Facility (SSRF, China), where data was collected. This work was supported by grants from the National Natural Science Foundation of China (81902063, 81871639, 92169109) and National Key R&D Program of China (2018YFA0507100, 2020YFA0707500). This work was supported by the Natural Science Program of Chongqing Science and Technology Commission (cstc2019jcyj-msxmX0135), Science and Technology Research pogram of Chongqing Municipal Education Commission (KJQN202000442) and CQMU Program for Youth Innovation in Future Medicine (W0073) to W.W. This work was supported by grants from Lingang Laboratory (LG202101-01-07), Science and Technology Commission of Shanghai Municipality (YDZX20213100001556 and 20XD1422900), Natural Science Foundation of Tianjin (18JCJQJC48000) and Shanghai Municipal Science and Technology Major Project (ZD2021CY001) to H.Y. This work was supported by Shanghai Frontiers Science Center for Biomacromolecules and Precision Medicine, ShanghaiTech University.

## Author contributions

H.Y. and W.W. conceived the project; W.C., H.Y. and W.W. designed the experiments; W.C., H.H., Z.L., and T.Z., cloned, expressed, purified and crystallized proteins; Y.D. and H.W. collected the diffraction data; W.C., Y.D., H.W. and X.J. solved the crystal structure; H.H., W.C., Y.D., W.S., D.S., X.J., W.W. and H.Y. analyzed and discussed the data; H.N., X.J., W.W. and H.Y. wrote the manuscript.

