## Supplemental Table 1, Figure S1 and S2 for "Crystal structure of monkeypox H1 phosphatase, an Antiviral Drug Target"

Extended data Table 1 Data collection and refinement statistics

| Property | Value |
|---|---|
| **Data collection** | |
| Space group | *I* 4 2 2 |
| Cell dimensions | |
| a, b, c (Å) | 62.48, 62.48, 170.494 |
| α, β, γ (°) | 90.00, 90.00, 90.00 |
| Resolution (Å) | 30.67-1.80 (1.86-1.80) |
| $I/\sigma I$ | 26.6 (4.4) |
| Redundancy | 16.1 (16.7) |
| Completeness (%) | 100.0 (100.0) |
| $R_{merge}$ | 7.4% (63.4%) |
| $CC_{1/2}$ | 99.9 (95.6) |
| **Refinement** | |
| Resolution (Å) | 30.67-1.80 (1.83-1.80) |
| No. reflections | 16132(1586) |
| $R_{work}$ / $R_{free}$ | 0.172/0.196 (0.225/0.275) |
| No. atoms | |
| Protein | 1385 |
| Ligand/ion | 5 |
| Water | 99 |
| B-factors | |
| Protein | 28.74 |
| Ligand/ion | 26.63 |
| Water | 36.90 |
| R.m.s. deviations | |
| Bond lengths (Å) | 0.008 |
| Bond angles (°) | 0.094 |

Statistics for the highest-resolution shell are shown in parentheses.

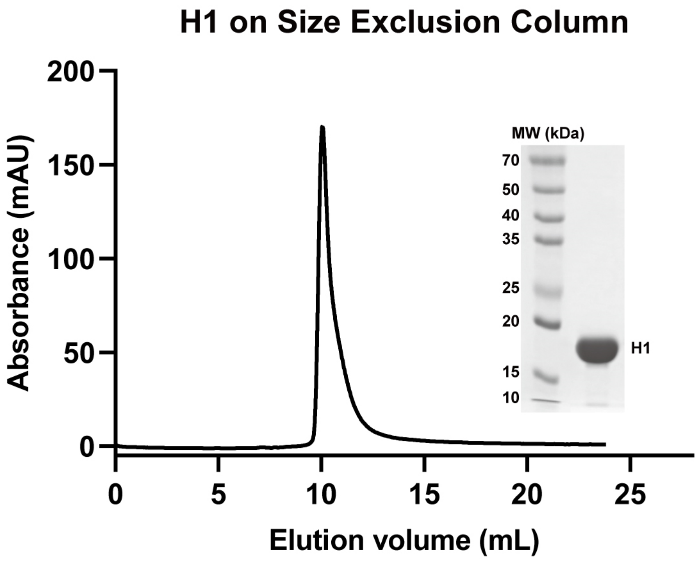


**Extended Data Fig. 1 | Size exclusion chromatography of monkeypox virus H1.** The curve shows H1 was eluted at 10 mL on a Superdex 75 increase 10/300 gl column (Cytiva, USA), which corresponds to a molecular weight of ~40 kDa. Since the molecular weight of H1 is ~18 kDa, the size exclusion chromatography suggests that H1 is a dimer in solution. SDS-PAGE analysis of the purified H1 is shown on the right.

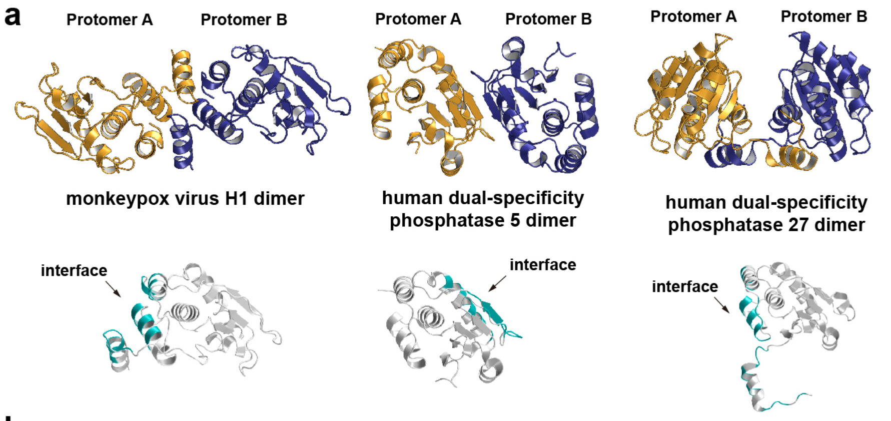


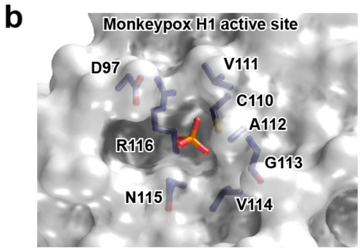


| Residues in active site of Monkeypox H1 | D97 | C110 | V111 | A112 | G113 | V114 | N115 | R116 |
|---|---|---|---|---|---|---|---|---|
| Corresponding residues in hDSP3 | D | C | R | E | G | Y | S | R |
| Corresponding residues in hDSP5 | D | S | E | A | G | I | S | R |
| Corresponding residues in hDSP10 | D | C | Q | A | G | V | S | R |
| Corresponding residues in hDSP27 | D | C | V | M | G | R | S | R |

**Extended Data Fig. 2 | Comparison between monkeypox H1 and human phosphatases with similar structure. a,** Monkeypox H1 and representative human phosphatases in the protein tyrosine phosphatase (PTP)/dual specificity phosphatase (DSP) family have different dimerization mode. The overall structures of dimeric monkeypox virus H1, human dual-specificity phosphatase 5 (hDSP5, PDB ID: 2G6Z) and human dual-specificity phosphatase 27 (hDSP27, PDB ID: 2Y96) are shown in the upper panel. The interface are in teal in the lower panel. **b**, the active site sequence is different among the phosphatases in the PTP/DSP family. The structure of the H1 active site is shown in the left panel. The residues at the active site of representative phosphatases are shown in the right panel. hDSP3: human dual-specificity phosphatase 3; hDSP10: human dual-specificity phosphatase 10.
